# Neutrophils prime unique transcriptional responses in intestinal organoids during infection with nontyphoidal *Salmonella enterica* serovars

**DOI:** 10.1101/2022.08.09.503428

**Authors:** Anna-Lisa E. Lawrence, Ryan P. Berger, David R. Hill, Sha Huang, Veda K. Yadagiri, Brooke Bons, Courtney Fields, Jason S. Knight, Christiane E. Wobus, Jason R. Spence, Vincent B. Young, Basel H. Abuaita, Mary X. O’Riordan

## Abstract

Nontyphoidal strains of *Salmonella enterica* are a major cause of foodborne illnesses and infection with these bacteria result in inflammatory gastroenteritis. Neutrophils are a dominant immune cell type found at the site of infection in *Salmonella-*infected individuals, but how they regulate infection outcome is not well understood. Here we used a co-culture model of primary human neutrophils and human intestinal organoids to probe the role of neutrophils during infection with two of the most prevalent *Salmonella* serovars: *Salmonella enterica* serovar Enteritidis and Typhimurium. Using a transcriptomics approach, we identified a dominant role for neutrophils in mounting differential immune responses including production of pro-inflammatory cytokines, chemokines, and antimicrobial peptides. We also identified specific gene sets that are induced by neutrophils in response to Enteritidis or Typhimurium infection. By comparing host responses to these serovars, we uncovered differential regulation of host metabolic pathways particularly induction of cholesterol biosynthetic pathways during Typhimurium infection and suppression of RNA metabolism during Enteritidis infection. Together these findings provide insight into the role of human neutrophils in modulating different host responses to pathogens that cause similar disease in humans.

**Importance:** Nontyphoidal serovars of *Salmonella enterica* are known to induce robust neutrophil recruitment in the gut during early stages of infection, but the specific role of neutrophils in regulating infection outcome of different serovars is poorly understood. Due to differences in human infection progression compared to small animal models, characterizing the role of neutrophils during infection has been challenging. Here we used a co-culture model of human intestinal organoids with human primary neutrophils to study the role of neutrophils during infection of human intestinal epithelium. Using a transcriptomics approach, we define neutrophil-dependent reprogramming of the host response to *Salmonella*, establishing a clear role in amplifying pro-inflammatory gene expression. Additionally, the host response driven by neutrophils differed between two similar nontyphoidal *Salmonella* serovars. These findings highlight the importance of building more physiological infection models to replicate human infection conditions to study host responses specific to individual pathogens.

## Introduction

Foodborne illnesses account for an estimated 48 million infections in the United States every year with 128,000 individuals needing to be hospitalized (1). One of the most common causes of foodborne disease is *Salmonella enterica*, which is responsible for an estimated 1.35 million infections in the United States each year (2). *Salmonella enterica* is comprised of over 2500 different serovars with *Salmonella enterica* serovar Typhimurium (STM) and Enteritidis (SE) among the most prevalent serovars globally. *Salmonella enterica* infects via the fecal-oral route and once it reaches the intestinal tract it stimulates a strong inflammatory response from the host leading to gastroenteritis and diarrheal disease (3). Although these symptoms are usually self-resolving, individuals with compromised immune systems or malnutrition can experience severe systemic, sometimes fatal, illness (4).

*Salmonella* pathogenesis is commonly studied *in vivo* using inbred mouse models. Notably, disease progression caused by *Salmonella* is often different in mice compared to humans, including the fact that mice rarely develop diarrhea during these infections (5). To better understand human infection, human-derived cells including human intestinal organoids (HIOs) have been used to define human-specific host responses to *Salmonella* (6–8). HIOs are derived from human pluripotent stem cells and self-organize to form a 3-dimensional polarized epithelium with differentiated epithelial cells and an underlying mesenchyme (9). Bacteria, including *Salmonella*, are able to replicate and stimulate robust inflammatory responses in HIOs (6–8, 10–12). Although the HIO model and other tissue culture models have been invaluable in revealing human-specific responses to *Salmonella* infection (6–8, 13–15), key features missing from these models are known to shape the outcome of infection, including immune cells.

Several immune cell types contribute to the control and resolution of *Salmonella* infections, however one of the earliest responders and the most abundant cell type found in *Salmonella*-infected individuals are polymorphonuclear leukocytes (PMNs), specifically neutrophils (16, 17). PMNs defend against bacterial infections through both cell-intrinsic and -extrinsic mechanisms - antimicrobial effectors like degradative proteases and ion chelators, production of reactive oxygen species and formation of sticky antimicrobial neutrophil extracellular traps (NETs) can directly kill bacteria (18,19)). PMNs also can influence surrounding cells and tissues, including the intestinal epithelium, changing the microenvironment via molecular oxygen depletion, regulating nutrient availability, and through production of inflammatory mediators (20, 21). How the interaction between intestinal epithelial cells and PMNs affect the outcome of bacterial infections is still poorly understood. To address this gap in knowledge, we generated a PMN-HIO model by co-culturing primary human PMNs with HIOs that were infected with *Salmonella enterica* serovar Enteritidis (SE) or Typhimurium (STM) by microinjection of bacteria into the lumen. Using this PMN-HIO model, we characterized how PMNs modulate intestinal epithelial host defenses during infection, compared to HIOs alone. We show here that the presence of PMNs elevates intestinal epithelial proinflammatory signaling, including production of cytokines, chemokines, and antimicrobial effectors. PMN-HIOs also distinguished between the two serovars through differential upregulation of metabolic pathways indicating that there are additional nuances in how PMN-HIOs respond to different pathogens.

## Results

### PMNs enhance HIO immune activation and other transcriptional responses during *Salmonella* infection

PMNs are potent drivers of inflammation, so we reasoned that PMNs would likely modulate the intestinal host response to *Salmonella* infection. Using the PMN-HIO model, we previously characterized the degree of PMN recruitment and effect on bacterial colonization during *Salmonella* infection (22). Here, to probe the contribution of PMNs to the global intestinal transcriptional responses during infection, HIOs and PMN-HIOs were microinjected with SE, STM or PBS and harvested at 8h post-infection (8hpi) for bulk RNA-sequencing (RNA-seq). Principal component analysis (PCA) was performed on normalized gene counts to determine whether PMNs drove separation between HIOs and PMN-HIOs and therefore changed the transcriptional profile during infection (Fig. 1A). While there was definitive segregation between infected HIOs in the presence and absence of PMNs, there was no clear separation between PBS control HIOs and PMN-HIOs, suggesting that PMNs change the transcriptional profile of the HIOs only during infection. To determine which biological pathways contributed most to the segregation between HIOs and PMN-HIOs, we extracted the loadings data from principal component 1 and subjected the top 50 genes to pathway enrichment analysis in Reactome to identify biological processes that these genes participate in (Fig. 1B). Approximately 75% of these genes belonged to the immune system, disease processes, and signal transduction suggesting a dominant role of immune activation for PMNs in the PMN-HIO model. Collectively, these data support that PMNs induce changes in HIO responses only during infection and that PMNs primarily induce enrichment of immune-related pathways.

**Fig 1.**
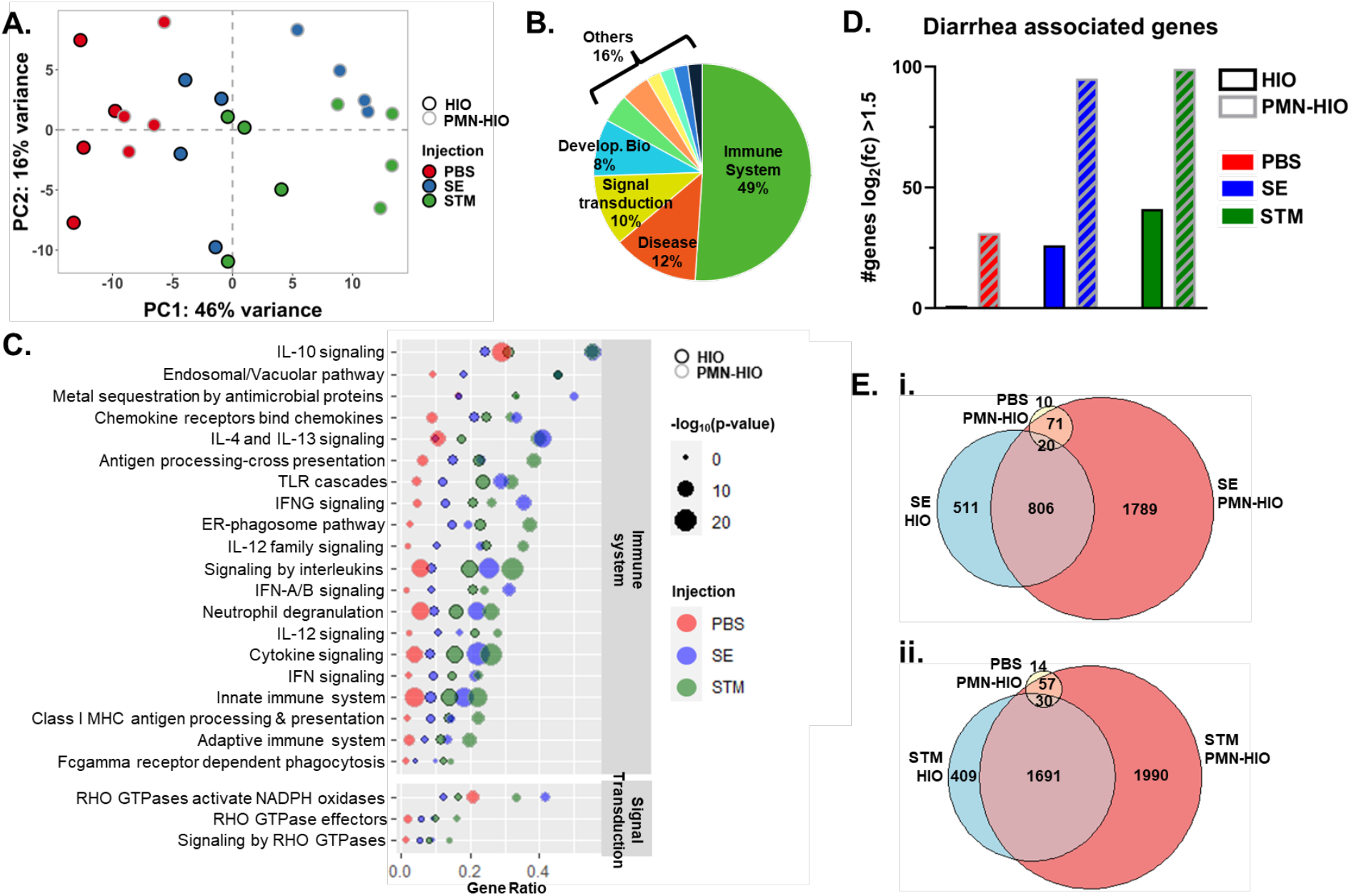
PMNs enhance HIO immune activation and other transcriptional responses during *Salmonella* infection. A.PCA plot of HIOs and PMN-HIOs infected with STM or SE with PBS as the mock control for 8h. B. Loadings data from PC1 was extracted from A to identify the top 50 genes driving separation of samples via PC1. Genes were then assessed for pathway enrichment using the Reactome database. Pie chart shows the percent of the top 50 pathways belonging to each of the Major Reactome pathway categories C. Dot plot assessing pathway enrichment of the top 50 identified pathways in (B) using all differentially expressed genes (p-adjusted value<0.05 compared to PBS-injected HIOs). Gene ratio is shown on the x-axis and the dot size corresponds to the −log10(p-value). HIO samples are outlined in black while PMN-HIOs are outlined in gray. PBS-injection (red), SE-injection (blue), STM-injection (green). D. Number of diarrhea associated genes with p-adjusted <0.05 and log2(fold change)>1.5 in each condition. Gene list for this analysis was used from (23). E. Venn diagram comparing differentially regulated genes with p-adjusted value <0.05 relative to PBS-injected HIOs for SE-injected HIOs, PBS-injected PMN-HIOs, and SE-injected PMN-HIOs (i.) and STM-injected HIOs, PBS-injected PMN-HIOs and STM-injected PMN-HIOs (ii.).

To further analyze which specific processes were induced by PMNs and how these pathways were enriched under various culture conditions, we performed pathway enrichment analysis of these top pathways with all genes that had a p-adjusted value <0.05 relative to control PBS-injected HIOs (Fig. 1C, and Table S1). Each pathway was analyzed for gene ratio (fraction of genes in a pathway that were significantly changed relative to total genes in that pathway) plotted on the x-axis, and the statistical significance, depicted as dot size, based on −log_10_(p-adjusted). As anticipated, immune system-related pathways were among the most significantly enriched pathways in response to infection (Fig. 1C). These included pathways belonging to processes involving signaling by interleukins, Toll-like receptor signaling, as well as PMN-specific pathways like neutrophil degranulation. We also detected enrichment of signal transduction pathways that also relate to PMN function. These pathways were related to Rho GTPase signaling. Activation of Rho GTPases has been shown in numerous studies to be critical in membrane remodeling during *Salmonella* infection and are targeted by secreted *Salmonella* effectors (23). Taken together, these results highlight the viability of the PMN-HIO model, demonstrating that neutrophils enhance the inflammatory tone of the HIO in an infection-specific manner.

One hallmark of infection with nontyphoidal *Salmonella* serovars is the development of inflammatory diarrhea. Since many of the top genes driving segregation between PMN-HIOs and HIOs were related to immune system processes, we assessed infected HIOs and PMN-HIOs for enrichment of genes associated with diarrhea (24). As predicted, the number of diarrhea-associated genes that were significantly changed increased with infection (Fig 1D). Consistent with our findings that PMNs upregulate immune pathways, the number of diarrhea-associated genes further increased in infected PMN-HIOs compared to infected HIOs. PMNs also appeared to increase the fold change of many genes that were changed in infected HIOs. These results suggest that PMNs not only amplify inflammatory processes during intestinal infection but may also promote induction of diarrhea.

To investigate how PMNs changed the HIO response during infection at the gene level, significant gene changes were calculated relative to PBS control HIOs and filtered for adjusted p-value <0.05 (Table S2). Venn diagrams were generated to compare gene changes during SE infection +/−PMNs or STM infection +/−PMNs (Fig. 1E). Although a substantial number of genes were changed during infection in both HIOs and PMN-HIOs, over 1700 additional genes were induced in SE-infected PMN-HIOs and over 1900 in STM-infected PMN-HIOs relative to HIOs alone. Importantly, there were very few genes induced in PBS control PMN-HIOs, confirming that adding PMNs to the HIOs alone does not trigger major changes in transcriptional programming, but the complex interaction between PMNs, HIOs and *Salmonella* drove a robust transcriptional response. Together these results demonstrate that co-culture with PMNs amplifies the HIO response to *Salmonella* infection including expression of genes that were not induced in infected HIOs alone, as well as enhanced enrichment of immune-related processes.

### PMNs elevate production of cytokines and chemokines in the PMN-HIOs

We recently reported that both SE and STM induce robust pro-inflammatory signaling in the HIO through transcriptional upregulation of cytokine and chemokine genes and downstream secretion of these effectors (6, 8). Because we observed a further increase in pathway enrichment of several pro-inflammatory pathways in the infected PMN-HIOs compared to infected HIOs, we examined the contribution of PMNs in changing expression and production of some of these pro-inflammatory mediators including cytokines and chemokines. Consistent with pathway enrichment results, PMN-HIOs increased expression of almost every cytokine and chemokine that was significantly changed during either SE or STM infection in the HIOs alone (Fig. 2A-2B). This elevated PMN-dependent response was primarily driven by infection, as there was little upregulation of these genes in PBS control PMN-HIOs. Of interest, in infected PMN-HIOs, we observed increased transcript levels of cytokines CSF-3, IL-6, IL-8 (Fig. 2A) and chemokines CXCL-10 and CCL-2 (Fig. 2B), all of which are essential for progression and resolution of intestinal inflammation (25–27). To assess whether these transcriptional changes were reflected at the protein level, supernatants from HIOs and PMN-HIOs were collected for ELISA to measure cytokine and chemokine output (Fig. 2C). Protein level analyses revealed similar patterns to the transcriptional results. Overall, the production of most cytokines and chemokines in infected PMN-HIOs was increased compared to infected HIOs or uninfected PMN-HIO controls. This included significant increases in IL-6, IL-8, CXCL-10, and CCL-2 production in infected PMN-HIOs compared to infected HIOs. However, some cytokines such as G-CSF (encoded by *CSF3*) or CXCL-2 did not significantly change with the addition of PMNs. While most other pro-inflammatory proteins correlated well with the transcript data, *CSF3* transcript was dramatically upregulated in infected PMN-HIOs, compared to infected HIOs, even though there was no difference in secreted protein levels. It is notable that although SE and STM induce similar degrees of *CSF3* transcript upregulation, protein levels of G-CSF in the supernatant are significantly lower in SE-infected PMN-HIOs compared to STM, suggesting that SE may regulate G-CSF post-transcriptionally. All together, we found that inflammatory signaling was elevated when PMNs are present in infected HIOs.

**Fig 2.**
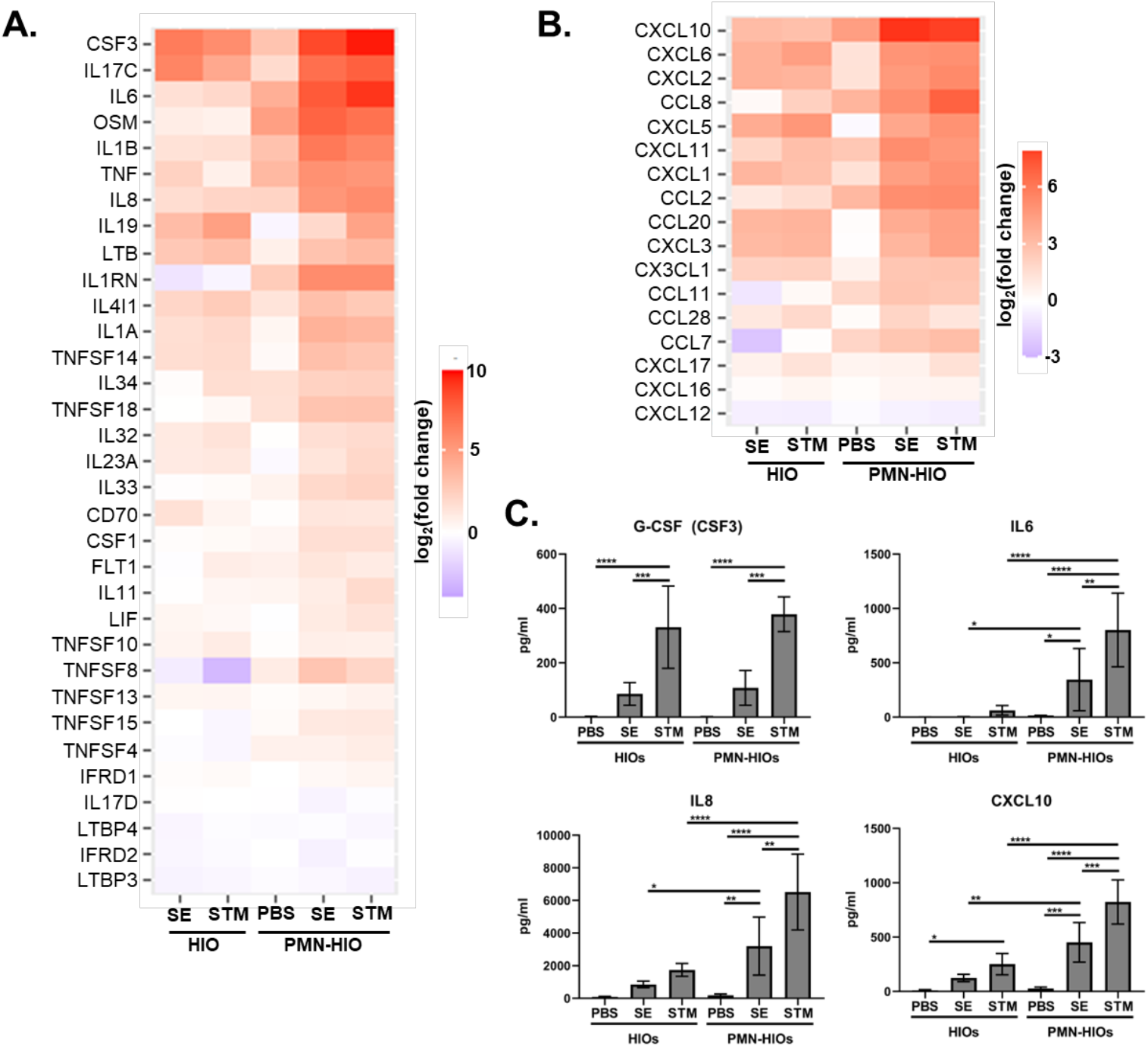
PMN-HIO co-culture amplifies production of cytokines and chemokines in infected HIOs. A-B. Gene expression data presented as log2(fold change) relative to PBS-injected HIOs for (A) cytokines and (B) chemokines. Genes that were significantly changed from PBS-injected HIOs in at least one condition with p-adjusted <0.05 are included. C. ELISA data from culture media sampled at 8hpi with 5 HIOs per well with n=4 replicates. Significance was determined by 2-way ANOVA where *p<0.05, **p<0.01, ***p<0.001, ****p<0.0001.

### HIO co-culture with PMNs strengthens the antimicrobial environment

Antimicrobial effectors such as defensins are a key component of the host defense in the intestine and our previous work demonstrated that antimicrobial peptides (AMPs) are strongly upregulated during infection in the HIO model (6, 8). Among the pathways enriched (Fig. 1C), was ‘metal sequestration by antimicrobial proteins’ suggesting that PMNs strongly affect the antimicrobial milieu in the HIOs. PMNs are known to have a wide arsenal of antimicrobial effectors, but specifically how PMNs affect the antimicrobial response during infection in complex environments such as in the PMN-HIO model has not been fully defined. In contrast to what we had predicted, PMNs did not dramatically enhance transcriptional upregulation of antimicrobial effectors above what was observed in infected HIOs. This included beta-defensins (DEFB4A/B), nutritional immunity effectors such as lipocalin (LCN2) or calprotectin (S100A8/9), and opsonins like SAA1/2 (Fig. 3A), which were all highly upregulated during infection in both HIOs and PMN-HIOs. Since PMNs represent approximately less than 5% of the cells in the PMN-HIO model (22), the contribution of PMN transcripts in the overall bulk-RNA-seq is likely minimal. However, analysis of culture supernatants via ELISA revealed that all these mediators were present at significantly higher levels in infected PMN-HIOs compared to HIOs (Fig. 3B). Some of these effectors were responsive to infection stimuli including beta-defensin (BD2, encoded by DEFB4A and DEFB4B) and elafin (encoded by PI3) as they were not present at high concentrations in the uninfected controls. In contrast, the nutritional immunity effectors calprotectin and lipocalin were constitutively produced by PMNs, whether HIOs were injected with PBS or *Salmonella*. Collectively, transcript and protein analyses of infected PMN-HIOs reveal that a small population of neutrophils can substantially enhance the antimicrobial environment.

**Fig 3.**
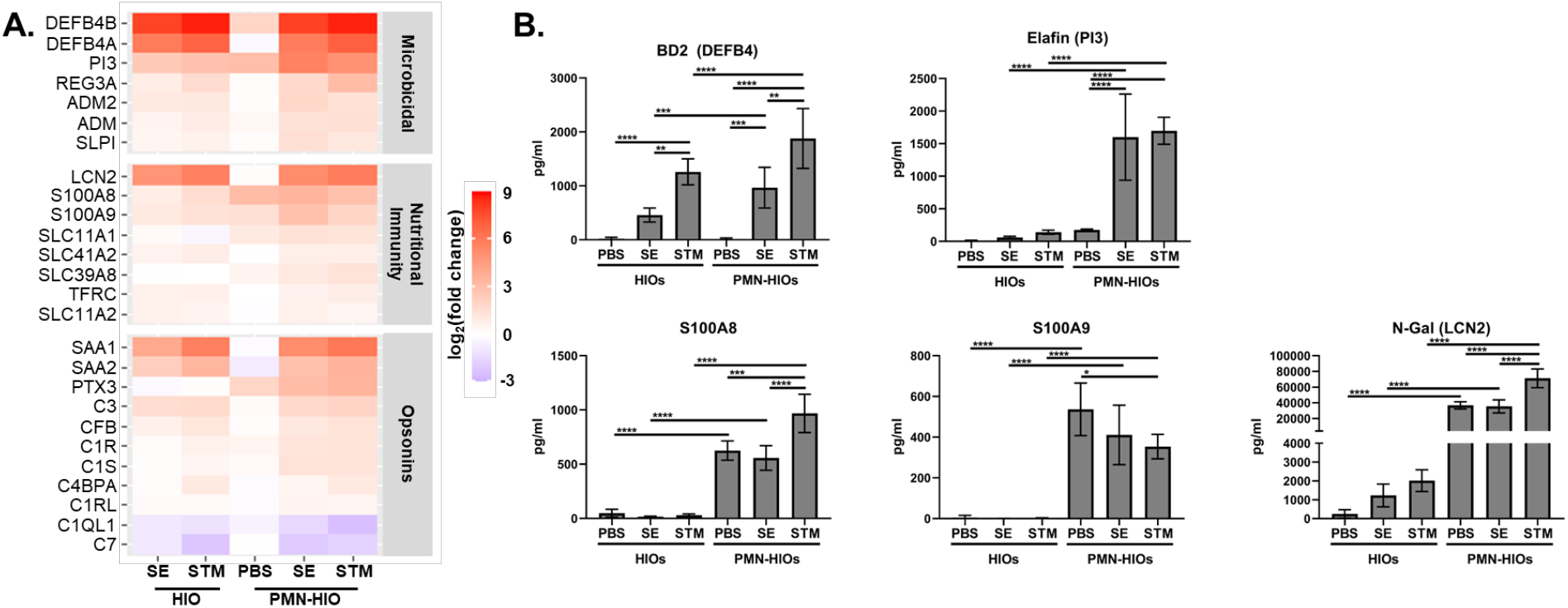
Presence of PMNs strengthens the antimicrobial environment. A. Gene expression data presented as log2(fold change) relative to PBS-injected HIOs for antimicrobial genes. Heatmap is divided into functional categories: Microbicidal, Nutritional Immunity, and Opsonins. Genes that were significantly changed from PBS-injected HIOs in at least one condition with p-adjusted <0.05 are included. B. ELISA data from culture media sampled at 8hpi with 5 HIOs per well with n=4 replicates. Significance was determined by 2-way ANOVA where *p<0.05, **p<0.01, ***p<0.001, ****p<0.0001.

### PMN-HIOs induce unique responses to two nontyphoidal *Salmonella* serovars

SE and STM share high levels of gene homology but accessory genes unique to SE contribute to its pathogenesis in a mouse model of infection (28–30). We also found that these serovars elicit unique responses by human intestinal epithelium (8). Consistent with this earlier finding, we observed clustering of SE-infected PMN-HIOs away from STM-infected PMN-HIOs via principal component analysis (PC2) (Fig. 1A). These data suggest that PMN-HIOs also mount distinct responses to these two serovars. To characterize these responses, significant genes from SE, STM, and PBS-injected PMN-HIOs were compared to determine the overlap in transcriptional responses to infection (Fig. 4A). Surprisingly, while there was a core set of host genes that were changed in response to both SE and STM, the majority of genes were uniquely induced by only one serovar with over 1000 genes uniquely changed during SE-infection of the PMN-HIOs and over 2000 genes uniquely changed during STM-infection. To better characterize the functional differences in these responses, the top 25 significant genes that were changed in either SE-infected or STM-infected PMN-HIOs were plotted and assessed for fold change relative to PBS-injected HIOs (Fig. 4B). The genes that were uniquely changed during SE-infection belonged to multiple biological processes including metabolism (PFKFB4 and HK2) although RNA modification-related genes were highly represented among the downregulated genes (ATIC, RRP12, NSUN2, and PUS1). More genes were downregulated in SE-infected PMN-HIOs compared to STM-infected PMN-HIOs where most genes that were uniquely responding to STM were upregulated. While these genes also fell into multiple categories, upregulation of cholesterol biosynthetic genes in STM-infected PMN-HIOs were highly prevalent in this heatmap (ACAT2, DHCR24, SCD, ACLY, and LSS). To further characterize which pathways were differentially induced during infection with the two serovars, pathway enrichment analysis on uniquely regulated genes identified in Fig. 4A was performed. To classify these pathways into different biological processes, each significantly enriched pathway was mapped back to the parent pathway in the Reactome database (Fig. 4C, Tables S3 and S4). Out of the 1012 genes that were uniquely induced in SE-infected PMN-HIOs, only 2 pathways were significantly enriched: ‘metabolism of RNA’ and ‘rRNA modification’. Most of the genes differentially expressed only during SE infected were downregulated (Fig. 4B). In contrast, 143 Reactome pathways were enriched in STM-infected PMN-HIOs representing a much more diverse set of biological processes. Cell cycle-related and signal transduction pathways were all highly represented among genes that were only induced in STM-infected PMN-HIOs, but we also noted that additional genes belonging to immune system pathways were uniquely induced in STM-infected PMN-HIOs leading to higher enrichment of these pathways. These findings suggest that while both serovars trigger different responses by PMN-HIOs, responses unique to SE infection largely relate to RNA metabolism while STM induces a broad range of host cell responses. Lastly, to compare pathway enrichment across all differentially expressed genes, pathways that were significant in either SE or STM infected PMN-HIOs were selected and sorted based on biggest difference in gene ratio between the two serovars (Fig. 4D). Multiple biological processes were differentially enriched when assessing the entire set of significant genes, including cell cycle, disease-related processes specifically relating to beta-catenin stability, immune system and metabolism related pathways. Pathways where SE exhibited greater gene ratios compared to STM-infected samples were mostly related to interferon signaling, although the ‘HIF1A stabilization’ pathway was also more enriched in SE-infected PMN-HIOs. Most of these pathways were also highly enriched during STM-infection with only a slight increase in gene ratio in the SE samples suggesting that both serovars trigger these pathways. In contrast, STM infection resulted in much stronger enrichment of beta-catenin related pathways and cholesterol biosynthesis pathways. Sub-pathways of cholesterol biosynthesis, namely biosynthesis via lathosterol and desmosterol had a gene ratio of 1 in STM-infected PMN-HIOs, meaning that all genes in these pathways were significantly changed during infection. Overall, our findings reveal differential responses between infection with SE and STM in the PMN-HIOs, including regulation of genes involved in RNA modification in SE-infected PMN-HIOs. Host responses to the two serovars were strikingly different in their stimulation of metabolic pathways, specifically highly enriched cholesterol biosynthetic pathways observed only in STM-infected PMN-HIOs.

**Fig 4.**
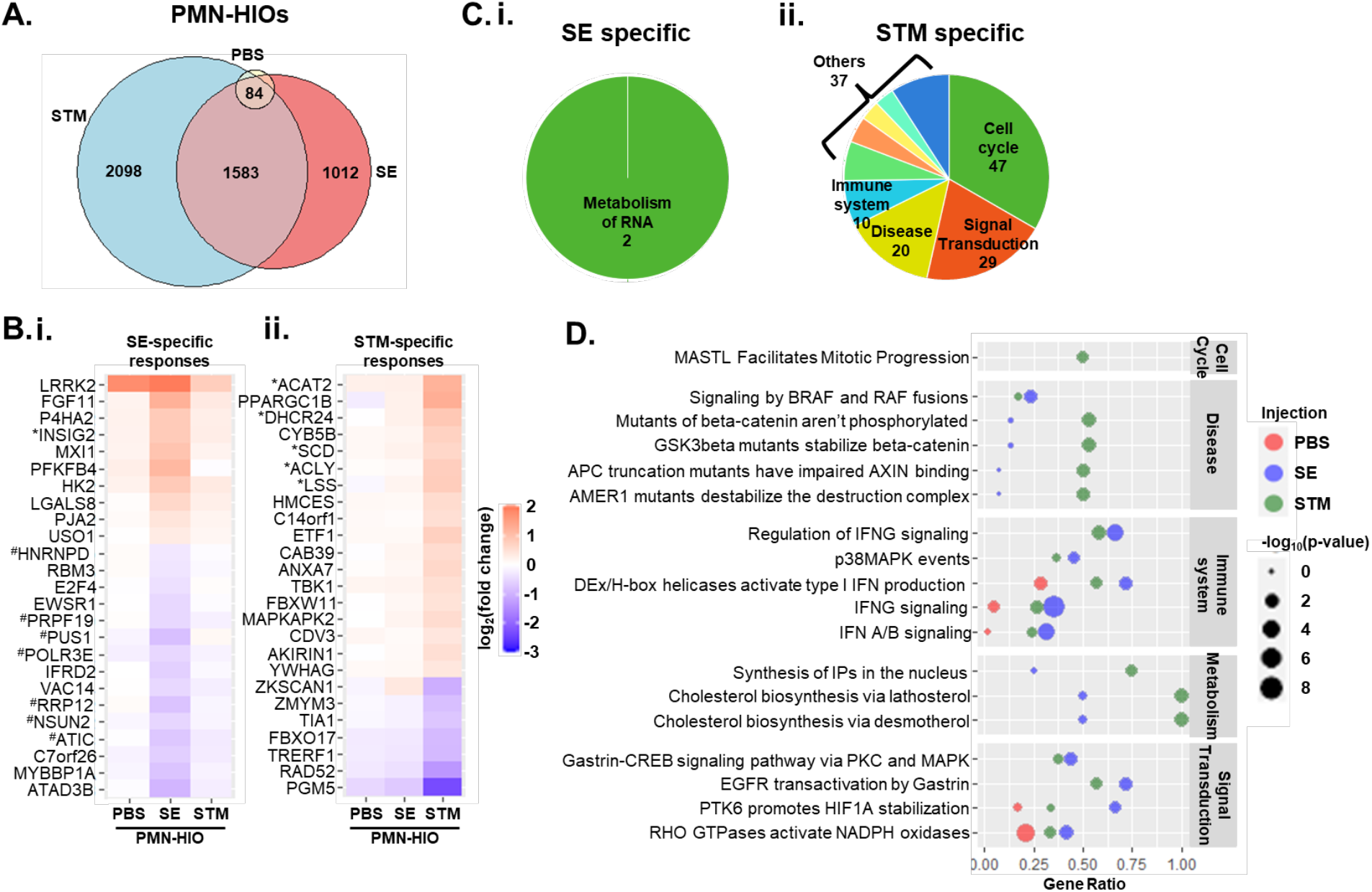
PMN-HIOs distinguish between SE and STM infection. A. Venn diagram comparing differentially regulated genes with p-adjusted value <0.05 relative to PBS-injected HIOs for STM-injected, SE-injected, or PBS-injected PMN-HIOs. B. Heatmap of unique differentially expressed genes. Top 25 genes based on adjusted p-value that were significant in only SE-injected PMN-HIOs (i.) or in STM-injected PMN-HIOs (ii.). Data are presented as log2(fold change) relative to PBS-injected HIOs. *Cholesterol biosynthetic genes, ^#^RNA modification genes. C. Pathway enrichment analysis of 1012 SE-induced genes (i.) or 2098 STM-induced genes identified in (A) from the Reactome database. Pie chart shows the number of each major Reactome pathway category represented by the pathway enrichment results out of 2 significant pathways in SE-infected PMN-HIOs or 143 significant pathways in STM-infected PMN-HIOs. D. Dot plot assessing pathway enrichment in PMN-HIOs of all differentially expressed genes (p-adjusted value<0.05 compared to PBS-injected HIOs). Top pathways were selected by the biggest difference in gene ratio between SE and STM-injected PMN-HIOs. Gene ratio is shown on the x-axis and the dot size corresponds to the −log10(p-value). PBS-injection (red), SE-injection (blue), STM-injection (green).

## Discussion

PMNs, particularly neutrophils, are early responders to inflammation and represent a dominant immune cell type recruited to the intestine during infection with nontyphoidal *Salmonella* serovars. Although it is appreciated that neutrophils are present during the early phases of *Salmonella* infection, how neutrophils contribute to the host response in a complex microenvironment like the intestine remains poorly understood. Here we used a transcriptomics approach in a PMN-HIO co-culture model to probe the role of PMNs during infection with two prevalent serovars of *Salmonella*: *Salmonella enterica* serovar Enteritidis and Typhimurium. We identified a dominant role for PMNs in enhancing inflammatory responses, including production of pro-inflammatory cytokines and chemokines as well as antimicrobial proteins during infection with both serovars. More broadly, PMNs acted on HIO cells to mount a more complex response to *Salmonella* as there were over 1000 genes in each infection condition that were uniquely and differentially expressed in the presence of PMNs. Lastly, we observed distinct responses between serovars; metabolic pathways were differentially regulated in the PMN-HIOs during SE and STM infection with robust enrichment of cholesterol biosynthesis genes during infection with STM, and downregulation of RNA modification-related genes during SE infection suggesting that PMNs can reprogram the environment in the presence of different pathogens.

Infection with nontyphoidal strains of *Salmonella enterica* induce robust inflammatory responses in the gut leading to gastroenteritis, and although neutrophils are known to be present at the site of infection, the specific contribution in tuning the inflammatory environment has been difficult to study in more complex animal models. We found via RNA-seq, as well as assessing secreted effectors, that the presence of PMNs during infection increased enrichment of cytokine signaling pathways, increased the upregulation of several cytokines that were already induced in infected HIOs alone, as well as led to a significant increase in levels of secreted cytokines and chemokines in the culture supernatants. This pattern was also conserved for antimicrobial effectors including a PMN-dependent upregulation in bactericidal and nutritional immunity related proteins. While a subset of these effectors are likely produced by PMNs such as calprotectin (encoded by S100A8/9) which is known to be highly produced in neutrophils, other effectors such as beta-defensins (encoded by DEFB4A/B) or elafin (encoded by PI3) are antimicrobial effectors known to be secreted by epithelial cells (31–33). This pattern would suggest that PMNs program the epithelium to enhance antimicrobial responses during infection. This pattern is supported by the prior finding that neutrophil elastase can induce production of elafin by epithelial cells (34). We also note a dramatic increase in the number of significant gene changes during infection in PMN-HIOs compared to HIOs, suggesting that PMNs likely stimulate a broad epithelial response to infection given the relatively low numbers of neutrophils in the PMN-HIOs. Further work may better elucidate the specific contribution of each cell type in the response to infection by performing single-cell RNA-seq, or other single cell approaches.

PMNs did not function to solely reprogram the inflammatory environment in response to *Salmonella* infection, but also affected other cellular processes that differentiated between infecting serovars. We found that over 3000 genes were differentially regulated during infection with SE and STM in the PMN-HIO model. One of the most striking differences we observed was transcriptional induction of cholesterol biosynthesis genes in STM infection of the PMN-HIOs, but not SE infection. In contrast, we noted that SE-infection uniquely induced upregulation of INSIG2, a negative regulator of cholesterol biosynthesis. There are numerous reports investigating the role of cholesterol during *Salmonella* infection including affecting invasion into epithelial cells, remodeling of the *Salmonella* containing vacuole (SCV), bacterial replication, and susceptibility to typhoid fever (35–43). Although there may be some cell type differences, the overarching pattern from these studies reveal that increased cholesterol generally enhances *Salmonella* infection. Our findings that SE does not induce upregulation of cholesterol biosynthetic genes is intriguing. It is possible that the two serovars have evolved to have a different dependence on cholesterol for productive infection. Alternatively, redistribution of existing cholesterol in the cells may be sufficient for SE. We previously reported that SE and STM exhibit different kinetics in their ability to invade and replicate within HIO epithelial cells, with SE invading at reduced frequency compared to STM (8). Our findings here point to cholesterol regulation as a potential mechanism that could contribute to differences in infection kinetics. Differential regulation of cholesterol biosynthesis as well as other serovar-specific responses warrant future study to better understand how these closely related serovars may be interacting with the intestinal epithelium in distinct ways.

The findings reported herein lay the groundwork for many future directions. In addition to PMN-dependent regulation of inflammatory responses and serovar-specific regulation of cholesterol biosynthesis genes, we also measured a shift in gene expression in response to *Salmonella* infection by using this co-culture model that was not observed in the HIO model alone. This included an increase in expression of diarrhea-associated genes. Modeling diarrheal diseases using mouse models has been difficult since mice rarely develop diarrhea in response to bacterial infections. Thus, our results open up an area for future study where the PMN-HIO co-culture model can be used to monitor human genes that are associated with diarrheal responses when infected with *Salmonella* or other microbial pathogens. Additionally, we show the feasibility of co-culturing HIOs with immune cells and in the future, this will allow us to add back components incrementally to model the intestinal environment and individually probe the role of immune cell subtypes in various pathological conditions. Further characterization of the HIO model and development of co-culture models of HIOs with immune cells will allow us to more closely study mechanisms governing the complex human-specific responses of the intestinal epithelium to bacterial infections.

## Acknowledgements

This work was supported by the NIH awards U19AI116482-01 (V.B.Y, J.R.S, C.E.W, and M.X.O) and R21AI13540 (M.X.O). A-L.E.L was supported by NIH T32 AI007528. We thank the Host Microbiome Initiative, the Comprehensive Cancer Center Immunology (supported by the NCI and NIH award P30CA046592) and Histology Cores and the DNA Sequencing Core at University of Michigan Medical School. We gratefully acknowledge the O’Riordan lab members for helpful discussions, and Dr. Roberto Cieza for assistance with data management.

## Author contributions

A-L.E.L and B.H.A performed the experiments and analyzed the data. A-L.E.L, M.X.O and B.H.A designed the experiments and wrote the manuscript. R.P.B and D.R.H assisted in RNA-seq analysis. S.H, V.K.Y, B.B and C.F assisted in HIO differentiation and culturing. J.R.S, V.B.Y, C.E.W and J.S.K assisted in experimental preparation and data interpretation.

## Materials and Methods

### Contact for reagent and resource sharing

All RNA sequences are deposited in the EMBL-EBI Arrayexpress database (E-MTAB-11089). Source code for RNA-seq analyses can be found at <u>aelawren/PMN-HIO-RNA-seq: R scripts for PMN-HIO RNA-seq analysis (github.com)</u>. Other reagents and resources can be obtained by directing requests to the Lead Contacts, Basel Abuaita and Mary O’Riordan.

### Experimental model and subject details

#### Human Intestinal Organoids (HIOs)

HIOs were generated by the *In Vivo* Animal and Human Studies Core at the University of Michigan Center for Gastrointestinal Research as previously described (9). Prior to experiments, HIOs were removed from the Matrigel, washed with DMEM:F12 media, and re-plated with 5 HIOs/well in 50μl of Matrigel (Corning) in ENR media ((DMEM:F12, 1X B27 supplement, 2mM L-glutamine, 100ng/ml EGF, 100ng/ml Noggin, 500ng/ml Rspondin1, and 15mM HEPES). Media was exchanged every 2-3 days for 7 days.

#### Human Polymorphonuclear Leukocytes (PMNs)

PMNs were isolated from blood of healthy human adult donors as previously described (44), according to the protocol approved by the University of Michigan Medical School (HUM00044257). Written consent was obtained from all donors. The purity of PMNs was assessed by flow cytometry using APC anti-CD16 and FITC anti-CD15 antibodies (Miltenyi Biotec), markers characteristic of human neutrophils.

#### Bacterial Growth and HIO Microinjection

*Salmonella enterica* serovar Typhimurium SL1344 (STM) and *Salmonella enterica* serovar Enteritidis P125109 (SE) were used throughout the manuscript. Bacteria were stored at −80°C in Luria-Bertani (LB, Fisher) medium containing 20% glycerol and cultured on LB agar plates. Individual colonies were grown overnight at 37ºC under static conditions in LB liquid broth. Bacteria were pelleted, washed and re-suspended in PBS. Bacterial inoculum was estimated based on OD_600_ and verified by plating serial dilutions on agar plates to determine colony forming units (CFU). Lumens of individual HIOs were microinjected with glass caliber needles with 1μl of PBS, SE or STM (10^5^ CFU/HIO) as previously described (6, 45, 46). HIOs were then washed with PBS and incubated for 2h at 37°C in ENR media. HIOs were treated with 100µg/ml gentamicin for 15 min to kill any bacteria outside the HIOs, then incubated in fresh medium +/−PMNs (5 × 10^5^ PMNs/5HIOs/well in a 24-well plate).

## Method Details

### Cytokine Analyses

For cytokine analysis, media from each well containing 5 HIOs/well were collected at 8hpi. Cytokines, chemokines and antimicrobial proteins were quantified by ELISA at the University of Michigan Cancer Center Immunology Core.

### RNA Sequencing and Analysis

Total RNA was isolated from 5 HIOs per group with a total of 4 replicates per condition using the mirVana miRNA Isolation Kit (Thermo Fisher). The quality of RNA was confirmed, ensuring the RNA integrity number (RIN)> 8.5, using the Agilent TapeStation system. cDNA libraries were prepared by the University of Michigan DNA Sequencing Core using the TruSeq Stranded mRNA Kit (Illumina) according to the manufacturer’s protocol. Libraries were sequenced on Illumina HiSeq 2500 platforms (single-end, 50 bp read length). All samples were sequenced at a depth of 10.5 million reads per sample or greater. Sequencing generated FASTQ files of transcript reads that were pseudoaligned to the human genome (GRCh38.p12) using kallisto software (47). Transcripts were converted to estimated gene counts using the tximport package (48) with gene annotation from Ensembl (49).

### Gene Expression and Pathway Enrichment Analysis

Differential expression analysis was performed using the DESeq2 package (50) with *P* values calculated by the Wald test and adjusted *P* values calculated using the Benjamani & Hochberg method (51). Pathway analysis was performed using the Reactome pathway database and pathway enrichment analysis in R using the ReactomePA software package (52).

### Quantification and Statistical Methods

RNA-seq data analysis was done using RStudio version 1.1.453. Plots were generated using ggplot2 (53) with data manipulation done using dplyr (54). Euler diagrams of gene changes were generated using the Eulerr package (55). Other data were analyzed using Graphpad Prism 9. Statistical differences were determined using statistical tests indicated in the fig. legends. The mean of at least 2 independent experiments were presented with error bars showing standard deviation (SD). *P* values of less than 0.05 were considered significant and designated by: \**P* < 0.05, \*\**P* < 0.01, \*\*\**P* < 0.001 and **** *P* < 0.0001.

## Supplemental information

**Table S1. Complete list of significant REACTOME pathways with p-value <0.05**. Genes with p-adjusted value <0.05 and log_2_(fold change) >0 relative to PBS-injected HIOs were selected for analysis.

**Table S2. Complete list of differentially expressed genes with p-adjusted value of <0.05 compared to PBS-injected HIOs**.

**Table S3. Complete list of significant REACTOME pathways from genes differentially expressed only in SE-infected PMN-HIOs**. Genes with p-adjusted value <0.05, log_2_(fold change) >0 relative to PBS-injected HIOs and p-adjusted value >0.05 in STM-infected PMN-HIOs were selected for analysis.

**Table S4. Complete list of significant REACTOME pathways from genes differentially expressed only in STM-infected PMN-HIOs**. Genes with p-adjusted value <0.05, log_2_(fold change) >0 relative to PBS-injected HIOs and p-adjusted value >0.05 in SE-infected PMN-HIOs were selected for analysis.

